# The dynamics and interactions of respiratory pathogen carriage among French pilgrims during the 2018 Hajj

**DOI:** 10.1101/791004

**Authors:** Van-Thuan Hoang, Thi-Loi Dao, Tran Duc Anh Ly, Khadidja Belhouchat, Kamel Larbi Chaht, Jean Gaudart, Bakridine Mmadi Mrenda, Tassadit Drali, Saber Yezli, Badriah Alotaibi, Pierre-Edouard Fournier, Didier Raoult, Philippe Parola, Vincent Pommier de Santi, Philippe Gautret

**Author notes:** Corresponding author: Philippe Gautret, VITROME, Institut Hospitalo-Universitaire Méditerranée Infection, 19-21 Boulevard Jean Moulin 13385 Marseille Cedex 05, France.

## Abstract

Respiratory tract infections are frequent among Hajj pilgrims. We conducted this study to describe the dynamics of the acquisition of respiratory pathogens, their potential interactions and risk factors for possible lower respiratory tract infection symptoms (LRTI) among French pilgrims during the 2018 Hajj. Pilgrims from Marseille who were participating in the Hajj were recruited. Each participant underwent four successive systematic nasopharyngeal swabs before and during their stay in Saudi Arabia. Carriage of the main respiratory pathogens was assessed by PCR. 121 pilgrims were included and 93.4% reported respiratory symptoms during the study period. Polymicrobial carriage was observed in 73.8% samples. The acquisition of rhinovirus, coronaviruses and S*taphylococcus aureus* occurred soon after arrival in Saudi Arabia and rates decreased gradually after days 5 and 6. In contrast, *Streptococcus pneumoniae* and *Klebsiella pneumoniae* carriage increased progressively until the end of the stay in Saudi Arabia. *Haemophilus influenzae* and *Moraxella catarrhalis* carriage increased starting around days 12 and 13, following an initial clearance. Influenza viruses were rarely isolated. We observed an independent positive mutual association between *S. aureus* and rhinovirus carriage and between *H. influenzae* and *M. catarrhalis* carriage. Dual carriage of *H. influenzae* and *M. catarrhalis* was strongly associated with *S. pneumoniae* carriage (OR = 6.22, 95%CI [2.04-19.01]). Finally, our model showed that *M. catarrhalis* carriage was negatively associated with *K. pneumoniae* carriage. Chronic respiratory disease was associated with symptoms of LRTI. *K. pneumoniae, M. catarrhalis-S. aureus* and *H. influenzae*-rhinovirus dual carriage was associated with LRTI symptoms. Our data suggest that RTIs at the Hajj are a result of complex interactions between a number of respiratory viruses and bacteria.

**Author summary:** Despite the recommendation to take individual preventive measures to prevent respiratory tract infections, these infections remain common among Hajj pilgrims. Respiratory pathogens acquired during the Hajj are usually studied individually, although in their natural environment they often compete or coexist with multiple microbial species. A better understanding of polymicrobial interactions in the nasopharynx among Hajj pilgrims is important. Our study describes the dynamics of the acquisition of respiratory pathogens and their potential interactions among pilgrims during the Hajj. We found that polymicrobial carriage was observed in most individuals and that some pathogens associated positively while other did not. Some pathogen associations correlated with symptoms of lower respiratory tract infections.

## Introduction

Each year, an increasing number of people travel to the Kingdom of Saudi Arabia (KSA) for the Hajj and Umrah pilgrimages, which attract around 10 million pilgrims annually from more than 180 countries. More than two million pilgrims from outside Saudi Arabia participated in the Hajj pilgrimages in 2017 [1]. Each year, about 2,000 pilgrims from Marseille, France, participate in the Hajj [2]. The event presents major challenges for public health, including inter-human transmission of infectious diseases, notably respiratory tract infections (RTIs), due the crowded conditions experienced by pilgrims [1]. In a recent study on morbidity and mortality among Indian Hajj pilgrims, infectious diseases represented 53% of outpatient diagnoses, with RTIs and gastroenteritis being the most common [3]. Between 69.8% and 86.8% of French pilgrims presented RTI symptoms during the Hajj [4]. A recent literature review suggested that etiology of RTIs at the Hajj is complex; several studies showed a significant acquisition of respiratory pathogens by pilgrims following participation in the Hajj in both symptomatic and asymptomatic individuals [5]. In a systematic review of 31 studies, Al-Tawfiq *et al*. showed that human rhinovirus (HRV) and influenza viruses were the most common viral respiratory pathogens isolated from ill Hajj pilgrims [6]. In addition, human non-MERS coronaviruses (HCoV) were also a common cause of RTIs during the event [7]. On the other hand, *Streptococcus pneumoniae*, *Haemophilus influenzae* and *Staphylococcus aureus* were shown to be the most commonly acquired respiratory bacteria at the Hajj [5].

RTIs are caused by the antagonistic and synergistic interactions between upper respiratory tract viruses and predominant bacterial pathogens [8]. Pathogens are usually studied individually, although in their natural environment they often compete or coexist with multiple microbial species. Similarly, the diagnosis of infections often proceeds via an approach which assumes a single-agent etiology [9]. Nevertheless, complex interactions occur between the different infectious microorganisms living in the same ecological niche and mixed infections are frequent [10].

A better understanding of polymicrobial interactions in the nasopharynx among Hajj pilgrims is important for many reasons. Carriage of more than one pathogen is common among Hajj pilgrims, whether or not they present with respiratory symptoms [7]. Colonisation is the initial step in the disease process [11]. Nasopharyngeal colonisation is likely to be a reservoir for respiratory pathogens resulting in interhuman transmission between pilgrims during close contact experienced during the Hajj ritual. Furthermore, antibiotic use or vaccines, which target specific pathogen species, may alter polymicrobial interactions in the nasopharynx and have unanticipated consequences [12, 13]. To our knowledge, the dynamics and interaction between the main respiratory pathogens acquired during the Hajj pilgrimage have not been specifically investigated, to date. Risk factors for possible lower respiratory tract infection (LRTI) symptoms at the Hajj have not been studied.

We conducted this study among French pilgrims during the 2018 Hajj, to describe the dynamics of the acquisition of respiratory pathogens and their potential interactions. In addition, we investigated risk factors for possible LRTI symptoms.

## Results

### Characteristics of study participants

The study included 121 pilgrims. The sex ratio of the population was 1:1.3 and the median age was 61 years with 58.7% of pilgrims aged 60 years and over. Most pilgrims were born in North Africa (66.9%) and sub-Saharan Africa (26.5%). There was a high prevalence of overweight (46.3%), obesity (28.1%), diabetes mellitus (25.6%) and hypertension (25.6%) and 13.2% participants reported that they suffered from chronic respiratory disease. In line with French recommendation, 88/121 pilgrims (72.7%) had an indication for vaccination against invasive pneumococcal disease (IPD) [14, 15] (Table 1).

**Table 1:** Characteristics of participants

| **Variables** | **N = 121** | **Prevalence (%)** |
| --- | --- | --- |
| **Gender** |  |  |
| Male | 52 | 43.0 |
| Female | 69 | 57.0 |
| **Age (years)** |  |  |
| Median |  | 61 |
| Interquartile |  | 56 - 66 |
| Min-Max |  | 26 - 83 |
| Age ≥60 | 71 | 58.7 |
| **Place of birth** |  |  |
| France | 8 | 6.6 |
| North Africa | 81 | 66.9 |
| Sub-Saharan Africa | 32 | 26.5 |
| **Comorbidities** |  |  |
| Diabetes mellitus | 31 | 25.6 |
| Hypertension | 31 | 25.6 |
| Chronic respiratory disease | 16 | 13.2 |
| Chronic heart disease | 13 | 10.7 |
| Chronic kidney disease | 3 | 2.5 |
| Immunodeficiency | 4 | 3.3 |
| **Indication for vaccination against IPD<sup>1</sup>** | **88** | **72.7** |
| **Smoking status** |  |  |
| Yes, current | 6 | 5.0 |
| Yes, stopped | 24 | 19.8 |
| Never | 91 | 75.2 |
| **BMI<sup>2</sup>** |  |  |
| Normal | 31 | 25.6 |
| Overweight | 56 | 46.3 |
| Obese | 34 | 28.1 |

<sup>1</sup>Age over or equal to 60 years, diabetes mellitus, chronic respiratory disease, chronic heart disease, chronic kidney disease and immunodeficiency
<sup>2</sup>Normal weight: BMI: 18.5 – 24.9, Overweight: BMI: 25.0 – 29.9, Obese: BMI $\geq 30$

A total of 37/121 (30.6%) pilgrims reported that they had been vaccinated against influenza in the past year. Only 17/88 (19.3%) pilgrims with an indication for IPD had been vaccinated against pneumococcal disease (PCV-13) in the past five years. Regarding non-pharmaceutical preventive measures, 49/121 (40.5%) pilgrims reported using face masks during the pilgrimage. Also, 67/121 (55.4%) and 70/121 (57.8%) pilgrims reported washing their hands more often than usual and using hand gel, respectively during the pilgrimage. Finally, 106/121 (87.6%) reported using disposable handkerchiefs during the Hajj.

### Clinical symptoms

A total of 113/121 (93.4%) pilgrims presented at least one respiratory symptom during their stay in KSA. A cough and rhinitis were the most frequent symptoms affecting 86.8% and 69.4% of participants. Over half of the pilgrims (59.5%) reported expectoration and 27.3% reported a dry cough. Voice failure was reported by 37.2%, fever by 27.3% and ILI by 20.7% of participants. Antibiotic use for RTIs was reported by 58.7% pilgrims. Only one (0.8%) pilgrim was hospitalised in KSA. Regarding possible LRTI symptoms, 5/121 (4.1%) participants reported a productive cough without nasal or throat symptoms. In addition, 9/121 (7.4%), 16/121 (13.2%) and 25/121 (20.7%) pilgrims presented febrile dyspnoea, dyspnea and a febrile productive cough, respectively. At total of 51/113 (45.1%) pilgrims with respiratory symptoms were still symptomatic at return. The mean time between arrival in KSA and the onset of symptoms was 8.7±4.6 days (range = 1-21 days) (data not shown). Most ill pilgrims presented the onset of respiratory symptoms when stationed at Mecca with a second minor wave in Mina (Fig 1).

**Fig 1.**
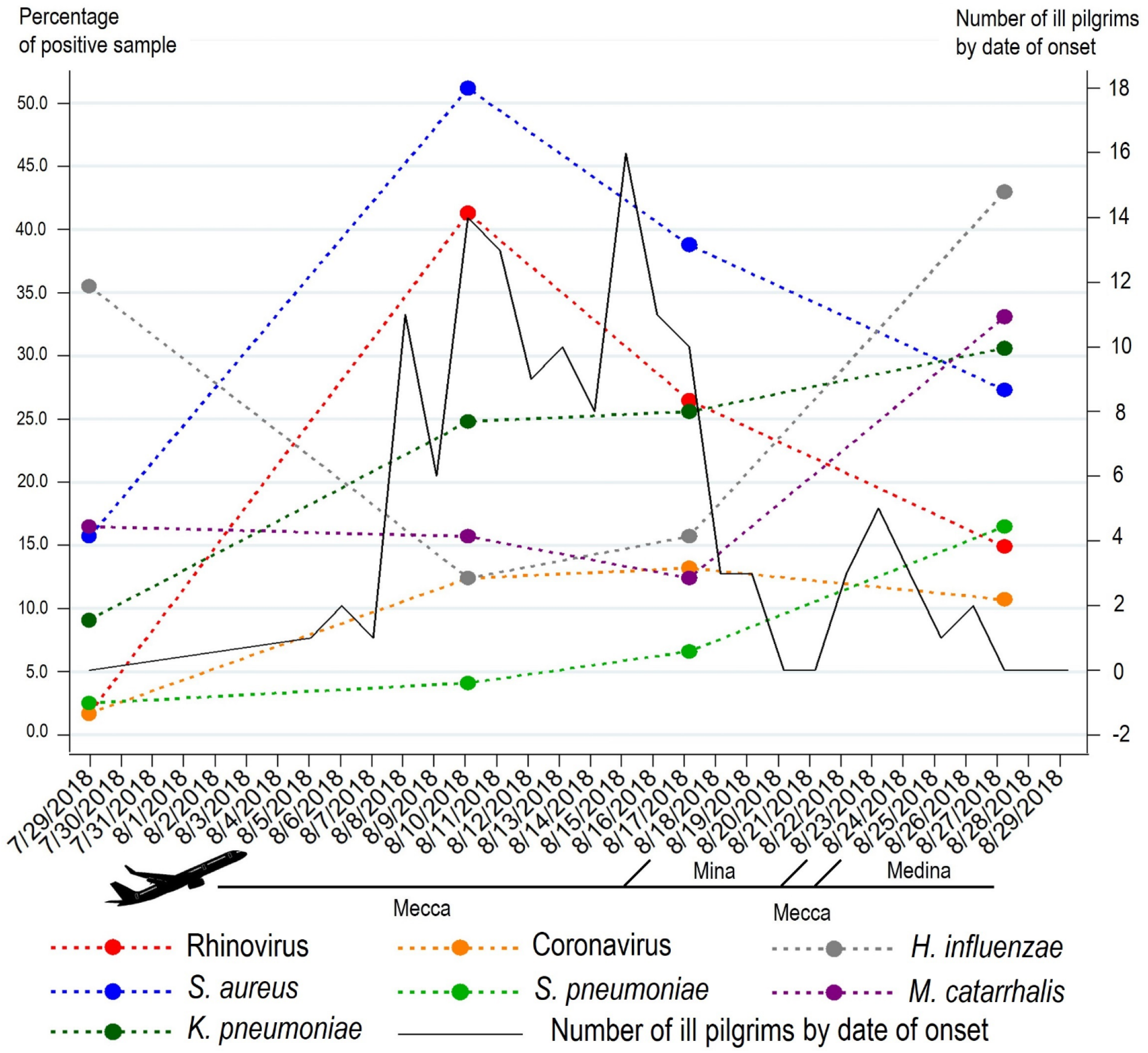
Dynamics of respiratory pathogens carriage during the 2018 Hajj and onset of respiratory symptoms among pilgrims

### Dynamics and interaction of respiratory pathogens carriage

Table 2 shows the prevalence of the carriage of respiratory pathogens according to sampling time and Figure 1 show the dynamics of most prevalent pathogens over the study period. Overall, 378/484 (78.1%) of all samples tested positive for at least one pathogen. *S. aureus* was the pathogen most frequently isolated with 33.3% of all samples testing positive. High positivity rates were also observed for *H. influenzae* (26.7%), *K. pneumoniae* (22.5%), HRV (21.1%) and *M. catarrhalis* (19.4%). Only 9.5% of the samples were positive for coronaviruses and 7.4% for *S. pneumoniae*. Very few samples tested positive for influenza viruses. Of the positive samples, the proportion which were positive for more than one pathogen was 210/378 (55.6%). A total of 138/378 (36.5%) samples were positive for two pathogens, 52/378 (13.8%) for three pathogens, 16/378 (4.2%) for four pathogens and 4/378 (1.1%) for five pathogens (data not shown).

**Table 2:** Prevalence of respiratory pathogens among pilgrims during the Hajj

|  | Pre-travel |  | Days 5 and 6 |  | Days 12 and 13 |  | Post-Hajj |  | Total |  |
| --- | --- | --- | --- | --- | --- | --- | --- | --- | --- | --- |
|  | N = 121 |  | N = 121 |  | N = 121 |  | N = 121 |  | N = 484 |  |
|  | n | % | n | % | n | % | n | % | n | % |
| **Respiratory viruses** |  |  |  |  |  |  |  |  |  |  |
| Influenza A | 0 | 0 | 0 | 0 | 0 | 0 | 3 | 2.5 | 3 | 0.6 |
| Influenza B | 0 | 0 | 2 | 1.7 | 1 | 0.8 | 1 | 0.8 | 4 | 0.8 |
| Rhinovirus | 2 | 1.7 | 50 | 41.3 | 32 | 26.5 | 18 | 14.9 | 102 | 21.1 |
| Coronavirus | 2 | 1.7 | 15 | 12.4 | 16 | 13.2 | 13 | 10.7 | 46 | 9.5 |
| **Respiratory bacteria** |  |  |  |  |  |  |  |  |  |  |
| *S. aureus* | 19 | 15.7 | 62 | 51.2 | 47 | 38.8 | 33 | 27.3 | 161 | 33.3 |
| *S. pneumoniae* | 3 | 2.5 | 5 | 4.1 | 8 | 6.6 | 20 | 16.5 | 36 | 7.4 |
| *H. influenzae* | 43 | 35.5 | 15 | 12.4 | 19 | 15.7 | 52 | 43.0 | 129 | 26.7 |
| *K. pneumoniae* | 11 | 9.1 | 30 | 24.8 | 31 | 25.6 | 37 | 30.6 | 109 | 22.5 |
| *M. catarrhalis* | 20 | 16.5 | 19 | 15.7 | 15 | 12.4 | 40 | 33.1 | 94 | 19.4 |
| **Virus-bacteria combination** |  |  |  |  |  |  |  |  |  |  |
| Rhinovirus-coronavirus | 0 | 0 | 5 | 4.1 | 1 | 0.8 | 2 | 1.7 | 8 | 1.7 |
| *S. aureus*-rhinovirus | 0 | 0 | 33 | 27.3 | 12 | 9.9 | 4 | 3.3 | 49 | 10.1 |
| *S. pneumoniae*-rhinovirus | 0 | 0 | 4 | 3.1 | 1 | 0.8 | 5 | 4.1 | 10 | 2.1 |
| *H. influenzae*-rhinovirus | 2 | 1.7 | 7 | 5.8 | 5 | 4.1 | 11 | 9.1 | 25 | 5.2 |
| *K. pneumoniae*-rhinovirus | 0 | 0 | 10 | 8.3 | 7 | 5.8 | 6 | 5.0 | 23 | 4.8 |
| *M. catarrhalis*-rhinovirus | 0 | 0 | 10 | 8.3 | 7 | 5.8 | 5 | 4.1 | 22 | 4.6 |
| *S. aureus*-coronavirus | 0 | 0 | 5 | 4.1 | 1 | 0.8 | 3 | 2.5 | 13 | 2.7 |
| *S. pneumoniae*-coronavirus | 0 | 0 | 1 | 0.8 | 2 | 1.7 | 2 | 1.7 | 4 | 0.8 |
| *H. influenzae*-coronavirus | 0 | 0 | 1 | 0.8 | 2 | 1.7 | 8 | 6.6 | 11 | 2.3 |
| *K. pneumoniae*-coronavirus | 0 | 0 | 5 | 4.1 | 3 | 2.5 | 2 | 1.7 | 10 | 2.1 |
| *M. catarrhalis*-coronavirus | 0 | 0 | 2 | 1.7 | 1 | 0.8 | 6 | 5.0 | 9 | 1.9 |
| **Bacteria combination** |  |  |  |  |  |  |  |  |  |  |
| *S. aureus*-*S. pneumoniae* | 2 | 1.7 | 2 | 1.7 | 1 | 0.8 | 7 | 5.8 | 12 | 2.5 |
| *S. aureus*-*H. influenzae* | 9 | 7.4 | 8 | 6.6 | 13 | 10.7 | 19 | 15.7 | 49 | 10.1 |
| *S. aureus*-*K. pneumoniae* | 3 | 2.5 | 13 | 10.7 | 12 | 9.9 | 6 | 5.0 | 34 | 7.0 |
| *S. aureus*-*M. catarrhalis* | 2 | 1.6 | 11 | 9.1 | 4 | 3.3 | 14 | 11.6 | 31 | 6.4 |
| *S. pneumoniae*-*H. influenzae* | 0 | 0 | 0 | 0 | 2 | 1.7 | 12 | 9.9 | 14 | 2.9 |
| *S. pneumoniae*-*K. pneumoniae* | 1 | 0.8 | 3 | 2.5 | 2 | 1.7 | 6 | 5.0 | 12 | 2.5 |
| *S. pneumoniae*-*M. catarrhalis* | 0 | 0 | 1 | 0.8 | 0 | 0 | 11 | 9.1 | 12 | 2.5 |
| *H. influenzae*-*K. pneumoniae* | 2 | 1.7 | 2 | 1.7 | 6 | 5.0 | 13 | 10.7 | 23 | 4.8 |
| *H. influenzae*-*M. catarrhalis* | 8 | 6.6 | 1 | 0.8 | 3 | 2.5 | 22 | 18.2 | 34 | 7.0 |
| *K. pneumoniae*-*M. catarrhalis* | 3 | 2.5 | 4 | 3.3 | 3 | 2.5 | 7 | 5.8 | 17 | 3.5 |

In pre-travel samples, virus carriage was very low with only a few participants testing positive for HRV and HCoV (<2%). Bacterial carriage was higher, notably for *H. influenzae* (35.5%) *M. catarrhalis* (16.5%) and *S. aureus* (15.7%). *K. pneumoniae* and *S. pneumoniae* carriage were relatively low (9.1% and 2.5%, respectively).

A dramatic increase in HRV carriage was observed on days 5 and 6 of the pilgrimage with prevalence 24 times higher than that of pre-travel. HRV carriage decreased progressively in subsequent samples but was still eight times higher in post-Hajj samples compared to pre-Hajj. A seven-fold increase of HCoV carriage was observed on days 5 and 6 that persisted on into days 12 and 13 of the pilgrimage and tended to slightly decrease in post-Hajj samples. Regarding bacteria, carriage of *S. aureus* increased by a factor of three on days 5 and 6 and decreased progressively in subsequent samples but was still double in post-Hajj samples compared to pre-Hajj. Interestingly, the carriage curves of HRV and *S. aureus* were strictly parallel.

*M. catarrhalis* carriage was about 12-16% in pre-travel, days 5 and 6 and 12 and 13 samples and increased to 33% in post-Hajj samples. *K. pneumoniae* carriage increased three-fold between pre-Hajj and days 5 and 6 samples and slightly increased in subsequent samples. *S. pneumoniae* carriage increased constantly overtime with a seven-fold increase in post-Hajj samples compared to pre-Hajj. Finally, *H. influenzae* carriage first decreased on days 5 and 6 and 12 and 13 by a factor 2.5 and then increased in post-Hajj samples to a carriage rate which was higher than that of pre-Hajj samples.

Table 3 shows the factors that were independently associated with the carriage of respiratory pathogens on 484 swabs from 121 pilgrims. A positive association was observed between males and carriage of HRV and *S. pneumoniae*. Chronic respiratory disease was also associated with *S. pneumoniae* carriage. Finally, the use of disposable handkerchiefs was associated with a decreased carriage of *H. influenzae*. Antibiotic intake ten days before each sampling was positively associated with HCoV and *K. pneumoniae* carriage.

**Table 3:**
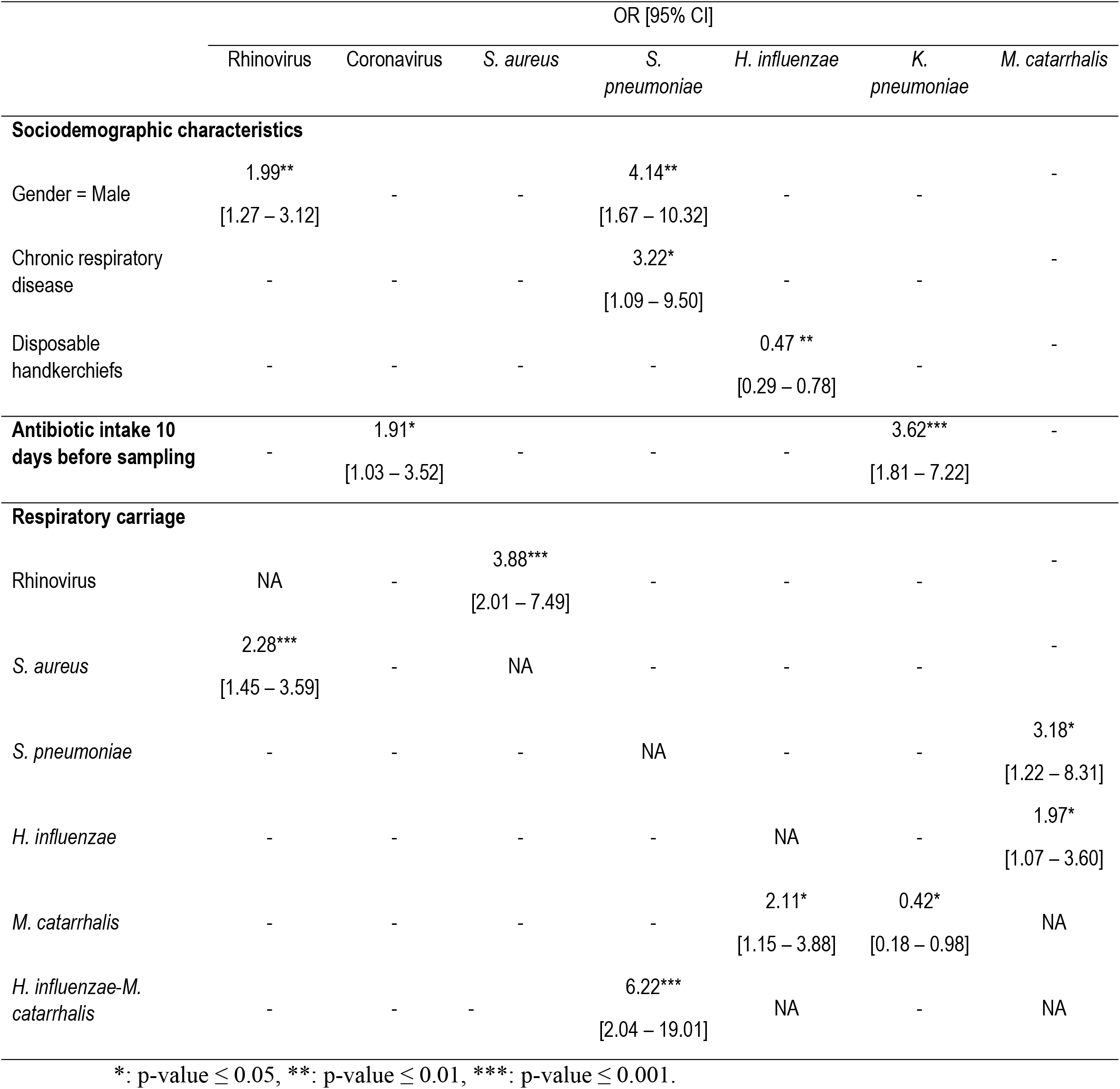
Risk factors for the carriage of respiratory pathogens among pilgrims during the Hajj (484 swabs from 121 pilgrims) (multivariable analysis)

|  | OR [95% CI] |  |  |  |  |  |  |
| --- | --- | --- | --- | --- | --- | --- | --- |
|  | Rhinovirus | Coronavirus | *S. aureus* | *S. pneumoniae* | *H. influenzae* | *K. pneumoniae* | *M. catarrhalis* |
| **Sociodemographic characteristics** |  |  |  |  |  |  |  |
| Gender = Male | 1.99**<br>[1.27 – 3.12] | - | - | 4.14**<br>[1.67 – 10.32] | - | - | - |
| Chronic respiratory disease | - | - | - | 3.22*<br>[1.09 – 9.50] | - | - | - |
| Disposable handkerchiefs | - | - | - | - | 0.47 **<br>[0.29 – 0.78] | - | - |
| **Antibiotic intake 10 days before sampling** | - | 1.91*<br>[1.03 – 3.52] | - | - | - | 3.62***<br>[1.81 – 7.22] | - |
| **Respiratory carriage** |  |  |  |  |  |  |  |
| Rhinovirus | NA | - | 3.88***<br>[2.01 – 7.49] | - | - | - | - |
| *S. aureus* | 2.28***<br>[1.45 – 3.59] | - | NA | - | - | - | - |
| *S. pneumoniae* | - | - | - | NA | - | - | 3.18*<br>[1.22 – 8.31] |
| *H. influenzae* | - | - | - | - | NA | - | 1.97*<br>[1.07 – 3.60] |
| *M. catarrhalis* | - | - | - | - | 2.11*<br>[1.15 – 3.88] | 0.42*<br>[0.18 – 0.98] | NA |
| *H. influenzae-M. catarrhalis* | - | - | - | 6.22***<br>[2.04 – 19.01] | NA | - | NA |

\*: p-value $\leq 0.05$ , \*\*: p-value $\leq 0.01$ , \*\*\*: p-value $\leq 0.001$ .

Regarding interactions between pathogens, we observed that HRV carriage and *S. aureus* carriage were mutually positively associated. The same applied to *H. influenzae* and *M. catarrhalis* carriage. Pilgrims carrying *S. pneumoniae* were more likely also to carry *M. catarrhalis*. Patients with a dual carriage of *H. influenzae* and *S. pneumoniae* were six times more likely also to be carrying *S. pneumoniae*. By contrast, *M. catarrhalis* carriage was associated with a reduced carriage of *K. pneumoniae*.

### Risk factors for possible lower respiratory tract infection among French pilgrims during the 2018 Hajj season

Table 4 shows the results of multivariable risk factor analysis for possible LRTI symptoms. Chronic respiratory disease was associated with all possible LRTI symptoms. Obesity was associated with dyspnea. Carriage of *K. pneumoniae* or *M. catarrhalis-S. aureus* or *H. influenzae*-rhinovirus combination was associated with a four-fold, 16-fold and eight-fold increase of dyspnoea prevalence, respectively. Finally, *M. catarrhalis-S. aureus* dual carriage was associated with a five-fold increase in the prevalence of febrile dyspnea (Table 4).

**Table 4:**
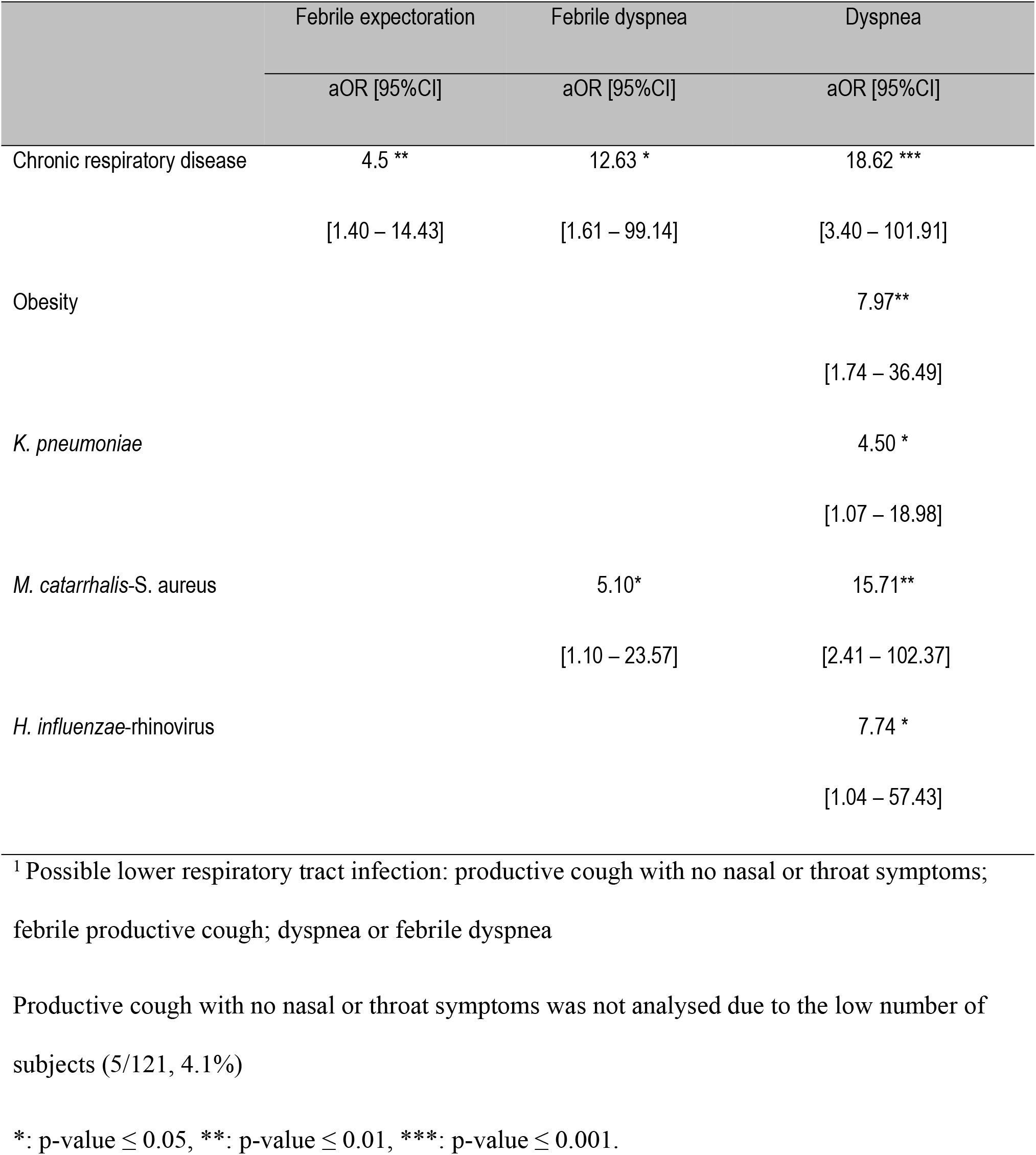
Risk factors for possible lower respiratory tract infection^1^ during the 2018 Hajj (multivariable analysis)

## Discussion

Despite the recommendation to take individual preventive measures to prevent RTIs [16, 17], these infections remain common among Hajj pilgrims. Overcrowding during the event is thought to increase the risk of the transmission of infectious diseases, but interaction between respiratory pathogens is probably one factor contributing towards the development of RTIs. To our knowledge, no study on respiratory microbiota alteration among pilgrims during the Hajj has been conducted. Our results about the occurrence of RTI symptoms are in line with previous results obtained regarding French pilgrims [18] and others [1, 19]. Notably, we observed that RTI symptoms occur soon after the pilgrims’ arrival in Mecca, with most symptoms starting between 4 to 13 days after arrival, corresponding to the period when pilgrims are stationed in Mecca hotels and are visiting the Grand Mosque daily, where highly crowded conditions are common [7]. We also confirmed that an overall increase in the carriage of respiratory viruses and bacteria can be seen when comparing pre-travel samples and post-Hajj samples, as previously documented [7, 12, 19–22]. Higher acquisition rates were observed for rhinovirus with a ninefold increase when comparing pre-travel to post-Hajj carriage and for *S. pneumoniae* with a seven-fold increase, but an increase was observed for all pathogens tested in this study. The unique design of our study with sequential systematic sampling at regular intervals allows for a better understanding of the dynamics of pathogen carriage during the pilgrimage.

Carriage rates of bacteria and viruses in this study are in line with those observed during recent studies conducted on French pilgrims and in pilgrims of other nationalities, using the same methods of detection [7, 19–22].

The acquisition of respiratory viruses and *S. aureus* occurred soon after arrival in Saudi Arabia and decreased gradually after days 5 and 6. By contrast, *S. pneumoniae* and *K. pneumoniae* carriage increased progressively until the end of the visit, *H. influenzae* and *M. catarrhalis* carriage increased later, after an initial clearance.

We hypothesise that the brutal acquisition of respiratory viruses upon arrival was the initial step that triggered subsequent changes in the relative abundance of resident bacteria [23] that were already present in the nasopharynx of pilgrims. The apparent simultaneity of viruses and *S. aureus* carriage increase and the initial wave of respiratory symptoms, suggests that this pathogen association was responsible for the RTIs that affected most pilgrims soon after arriving in Mecca. The subsequent increase in resident bacteria that occurred during the second half of pilgrims’ stays in Saudi Arabia appears to be contemporaneous with a second wave of respiratory symptoms, suggesting that these RTIs were of bacterial origin.

Regarding interaction between respiratory pathogens, we observed a very clear pattern of positive association between the carriage of *S. aureus* and rhinovirus with acquisition curves which were strictly parallel. Furthermore, an independent positive mutual association between the carriage of the two pathogens was evidenced in our multivariate model. Several studies revealed a positive interaction between natural or experimental rhinovirus infection and *S. aureus* nasal carriers [24–28]. These studies also underlined that rhinovirus infection may facilitate the propagation of *S. aureus* from staphylococcal carriers to the environment and the transmission of the bacterium between humans. Among healthy persons who were experimentally infected by rhinovirus, the relative abundance of *S. aureus* first increased and then returned to its baseline level after the rhinovirus infection was cleared [29]. These results suggest that changes in the composition of the respiratory microbiota following rhinovirus infection may play a role in the development of bacterial superinfection. Morgene *et al*. proposed several potential mechanisms through which rhinovirus may increase bacterial infection [30]. Rhinovirus infection promotes pro-inflammatory cytokines and IFN production mainly through the activation of NFκB. In rhinovirus infected cells, the adherence of *S. aureus* was significantly higher compared to uninfected cells. The inflammation due to rhinovirus infection also increased cellular patterns that facilitate the adhesion and internalisation of *S. aureus* within host cells [30].

We also observed a parallel increase of *H. influenzae* and *M. catarrhalis* carriage in days 12 and 13 and post-Hajj samples. An independent positive mutual association between the carriage of the two pathogens was evidenced in our multivariate model. Dual carriage of *H. influenzae* and *M. catarrhalis* strongly associated with *S. pneumoniae* carriage which in turns associated with *M catarrhalis* carriage. These results are consistent with those of several studies conducted among children with upper RTIs [31–33]. In these studies, the competitive interaction between *S. pneumoniae* and *H. influenzae* was dependent on neutrophils and complement. The additional carriage of *M. catarrhalis* might alter the competitive balance between *H. influenzae* and *S. pneumoniae* [32]. Co-colonisation of *S. pneumoniae* or *H. influenzae* with *M. catarrhalis* associating with increased risk of otitis media has been documented [34]. Using in vivo models, mixed species biofilms play a role in increasing the persistence of ear disease [35]. Other proposed mechanisms for positive associations between bacterial species include interspecies quorum sensing and passive antimicrobial resistance, which have been observed in experimental models of otitis media [36].

Finally, our model showed that *M. catarrhalis* carriage was negatively associated with *K. pneumoniae* carriage which, to our knowledge, has not previously been published.

Additionally, we found that the male gender was independently associated with an increase in rhinovirus and *S. pneumoniae* carriage. We have no explanation for this unexpected observation. The carriage of *S. pneumoniae* was higher among pilgrims with chronic respiratory disease which support the current French recommendations that vaccination against IPD be proposed to at-risk pilgrims [37]. In one of our recent studies, pilgrims who were vaccinated against IPD were seven time less likely to harbour *S. pneumoniae* after the Hajj compared to unvaccinated pilgrims [20]. In this study, the use of disposable handkerchiefs was associated with a significant decrease in *H. influenzae* carriage. Non-pharmaceutical individual preventive measures such as cough etiquette, hand hygiene, use of a face mask, disinfectant gel and disposable handkerchiefs are recommended for Hajj pilgrims [17]. Nevertheless, the effectiveness of these measures has been poorly investigated and available results are contradictory [17]. The apparent association between antibiotic use and HCoV and *K. pneumoniae* carriage warrants further investigation to better explore this unexpected observation.

We also confirm that chronic respiratory disease is a risk factor for LRTI. We also evidenced the role of respiratory bacteria including *K. pneumoniae* and *M. catarrhalis-S. aureus* association and *H. infuenzae*-rhinovirus association in the occurrence of possible LRTI symptoms. This reinforces the need for antibiotic use in case of LRTI symptoms [18].

Our study has some limitations. The study was conducted among French pilgrims only with a relatively small sample size and cannot be generalised to all pilgrims. qPCR used to detect respiratory pathogens does not distinguish between dead and living micro-organisms. Only a small number of respiratory pathogens were investigated. Respiratory bacteria serotypes were not investigated. Influenza viruses were not included in the model due to low carriage rates. Nevertheless, our study is the first study on the dynamics of and interaction between the respiratory pathogens that are most frequently isolated among Hajj pilgrims. Our data suggest that RTIs at the Hajj are a result of complex interactions between a number of respiratory viruses and bacteria. Further studies aimed at studying the respiratory microbiota with tools allowing the identification of larger numbers of pathogens will be necessary to better elucidate these ecological changes and their potential role in the occurrence of respiratory symptoms.

## Materials and methods

### Participants and study design (Fig 2)

**Fig 2.**
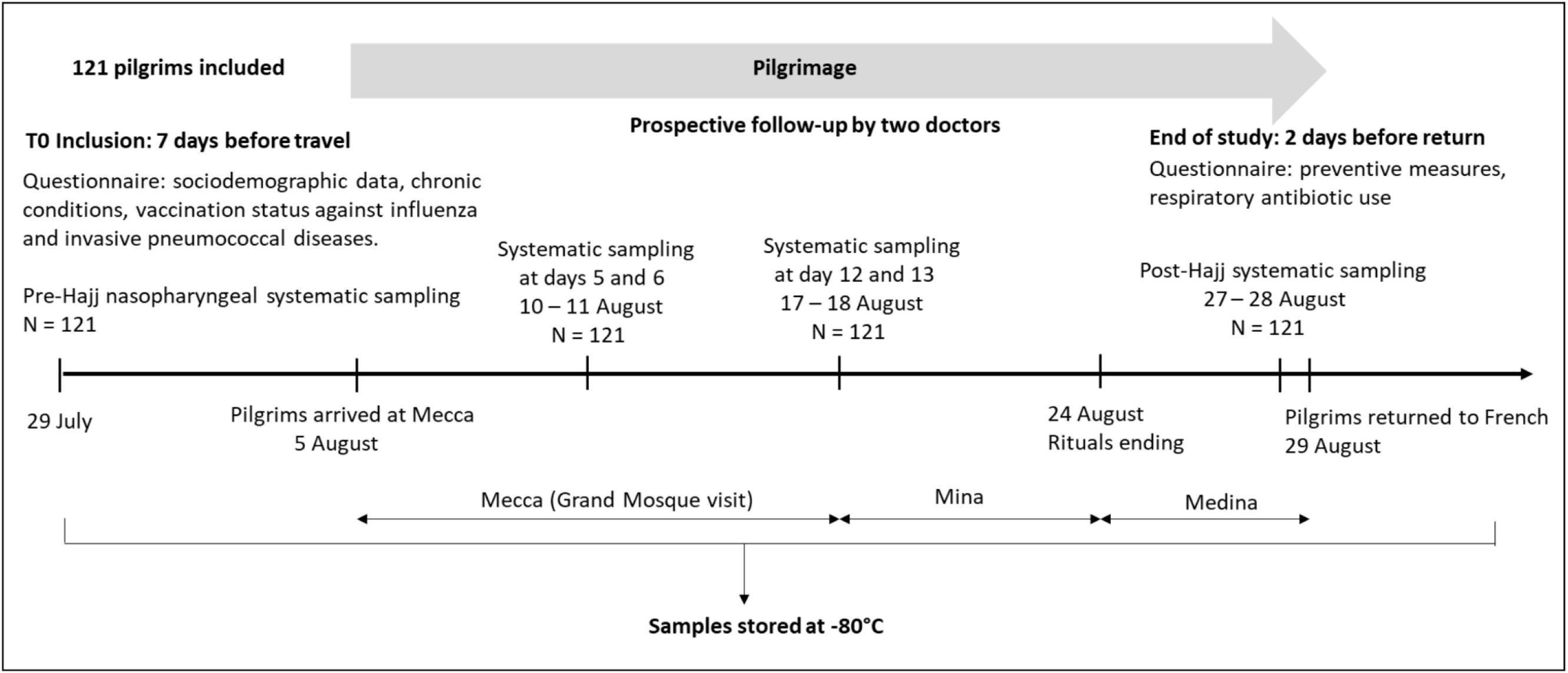
Study design of cohort survey among 121 French pilgrims in 2018 Hajj season

Pilgrims traveling to Mecca, Saudi Arabia during the 2018 Hajj from Marseille, France, were recruited through a private specialised travel agency. Potential adult participants were invited to participate in the study. They were included and followed-up by two bilingual (Arabic and French) medical doctors who travelled with the group. All participants departed to KSA on the same date, were housed in the same accommodation during their stay and performed the rituals together. Upon inclusion, before departing from France, pilgrims were interviewed using a standardised pre-Hajj questionnaire that collected information about demographic characteristics, medical history and immunisation status. Pilgrims were considered to have been immunised against influenza when they had been vaccinated within the past year and until before 10 days of the date of travel. Pilgrims were considered to be immune to IPD when they had been vaccinated with the 13-valent conjugate pneumococcal vaccine (PCV-13) in the past five years [14, 15]. A post-Hajj questionnaire was completed two days before the pilgrims’ return to France. Clinical data, antibiotic intake and information on compliance with face masks use as well as hand washing, the use of hand gel disinfectant and disposable handkerchiefs was collected. To evaluate the dynamic and interaction of respiratory pathogens during the Hajj, all pilgrims underwent four successive systematic nasopharyngeal swabs at different times: pre-travel, five to six days post arrival, 12 to 13 days post arrival and just prior to leaving KSA (post-Hajj). The Hajj rituals took place from 19–24 August. Influenza-like illness (ILI) was defined as the presence of cough, sore throat and subjective fever [38]. Possible LRTI was defined by presence of productive cough without nasal or throat symptoms; febrile productive cough; dyspnea or febrile dyspnea [18]. Based on the WHO classification, “underweight” was defined as having a body mass index (BMI) below 18.5, “normal” corresponded to a BMI between 18.5 and 25, “overweight” corresponded to a BMI ≥25, and “obese” referred to those with a BMI ≥30 [39].

#### Respiratory specimen

Nasopharyngeal swabs were obtained from each pilgrim, transferred to Sigma-Virocult® medium and stored at −80°C until processing.

### Identification of respiratory pathogens

The RNA and DNA were extracted from the samples using the EZ1 Advanced XL (Qiagen, Hilden, German) with the Virus Mini Kit v2.0 (Qiagen) according to the manufacturer’s recommendations. All quantitative real-time PCR were performed using a C1000 Touch™ Thermal Cycle (Bio-Rad, Hercules, CA, USA).

One-step simplex real-time quantitative RT-PCR amplifications were performed using Multiplex RNA Virus Master Kit (Roche Diagnostics, France) for influenza A, influenza B, HRV and internal controls MS2 phage [40]. HCoV was detected by one-step duplex quantitative RT-PCR amplifications of HCoV/ HPIV-R Gene Kit (REF: 71-045, BioMérieux, Marcy l’Etoile, France), according to the manufacturer’s recommendations.

Real-time PCR amplifications were carried out using LightCycler^®^ 480 Probes Master kit (Roche diagnostics, France) according to the manufacturer’s recommendations. The SHD gene of *H. influenzae*, phoE gene of *Klebsiella pneumoniae*, nucA gene of *S. aureus*, lytA CDC gene of *S. pneumoniae* and copB gene of *Moraxella catarrhalis* were amplified with internal DNA extraction controls TISS, as previously described [19, 41].

Negative controls (PCR mix) and positive controls (DNA from bacterial strain or RNA from viral strain) were included in each run. Positive results of bacteria or virus amplification were defined as those with a cycle threshold (CT) value ≤35.

Pilgrims were considered to be positive for respiratory pathogens during the Hajj if they were positive at the days 5 and 6 and/or days 12 and 13 sample.

## Statistical analysis

STATA software version 14.2 (Copyright 1985-2015 StataCorp LLC, http://www.stata.com) was used for statistical analysis.

The main outcomes of interest were the relationships between respiratory pathogens among pilgrims during the Hajj. We evaluated the carriage of HRV, HCoV, *S. aureus, S. pneumoniae, H. influenzae, K. pneumoniae* and *M. catarrhalis* using logistic mixed models. Because each pilgrim provided four successive samples, we used a repeated measures design to take into account the variability of series samples from each pilgrim. To evaluate the effect of covariates on each respiratory pathogen carriage, we modelled carriage of HRV, HCoV, *S. aureus, S. pneumoniae, H. influenzae, K. pneumoniae* and *M. catarrhalis* separately. We did not separately model the outcome of carriage of influenza A and B viruses because of the low prevalence of these viruses. Only the variables with a prevalence ≥5.0% were considered for statistical analysis. Unadjusted associations between respiratory pathogen carriage with multiples factors: sociodemographic characteristics (gender, ≥60 years), chronic respiratory disease, BMI classification, smoking status; individual preventive measures (vaccination against influenza, vaccination against IPD, use of a face mask, hand washing, disinfectant gel and disposable handkerchiefs); antibiotic intake 10 days before each sample; respiratory virus or bacteria and dual carriage were analysed by univariable analysis. Variables with p values <0.2 in the univariable analysis were included in the multivariable analysis. A mixed model with the subject being random effect was used to estimate the relationships between respiratory pathogens and to take into account the repeated measures for pathogen carriage for each subject.

Regarding risk factors for LRTI, the outcome was possible LRTI symptoms reported during the Hajj. The independent factors were sociodemographic characteristics (gender, ≥60 years), chronic respiratory disease, smoking status, BMI classification; vaccination against influenza, vaccination against IPD, respiratory virus or bacteria and dual carriage during the Hajj. Unadjusted associations between multiple factors and possible LRTI symptoms were examined using univariable analysis. Variables with p values <0.2 in the univariable analysis were included in the multivariable analysis. A logistical regression model was used to estimate factors’ adjusted odds ratios regarding possible LRTI.

The results were presented by odds ratio (OR) with a 95% confidence interval (95%CI). Results with a p value ≤0.05 was considered to be statistically significant.

## Ethics Statement

The protocol was approved by the Aix-Marseille University institutional review board (23 July 2013; reference No. 2013-A00961-44).

The study was performed according to the good clinical practices recommended by the Declaration of Helsinki and its amendments.

All participants provided their written informed consent.

## References

1. Memish ZA, Steffen R, White P, Dar O, Azhar EI, Sharma A, et al. Mass gatherings medicine: public health issues arising from mass gathering religious and sporting events. Lancet. 2019;393:2073–84. doi: 10.1016/S0140-6736(19)30501-X.

2. Gautret P, Bauge M, Simon F, Benkouiten S, Parola P, Brouqui P. Travel reported by pilgrims from Marseille, France before and after the 2010 Hajj. J Travel Med. 2012;19:130–2. doi: 10.1111/j.1708-8305.2011.00584.x

3. Khan ID, Khan SA, Asima B, Hussaini SB, Zakiuddin M, Faisal FA. Morbidity and mortality amongst Indian Hajj pilgrims: A 3-year experience of Indian Hajj medical mission in mass-gathering medicine J Infect Public Health. 2018;11:165–70. doi: 10.1016/j.jiph.2017.06.004

4. Gautret P, Benkouiten S, Griffiths K, Sridhar S. The inevitable Hajj cough: Surveillance data in French pilgrims, 2012-2014. Travel Med Infect Dis. 2015;13:485–9. doi: 10.1016/j.tmaid.2015.09.008.

5. Hoang VT, Gautret P. Infectious Diseases and Mass Gatherings. Curr Infect Dis Rep. 2018;20:44. doi: 10.1007/s11908-018-0650-9.

6. AlTawfq JA, Benkouiten S, Memish ZA. A systematic review of emerging respiratory viruses at the Hajj and possible coinfection with Streptococcus pneumoniae. Travel Med Infect Dis 2018;23: 6–13

7. Hoang VT, Sow D, Dogue F, Edouard S, Drali T, Yezli S, et al. Acquisition of respiratory viruses and presence of respiratory symptoms in French pilgrims during the 2016 Hajj: A prospective cohort study. Travel Med Infect Dis. 2019. pii: S1477–8939(19)30038-9. doi: 10.1016/j.tmaid.2019.03.003.

8. van den Bergh MR, Biesbroek G, Rossen JW, de Steenhuijsen Piters WA, Bosch AA, van Gils EJ, et al. Associations between pathogens in the upper respiratory tract of young children: interplay between viruses and bacteria. PLoS One. 2012;7:e47711. doi: 10.1371/journal.pone.0047711.

9. Brunstein JD, Cline CL, McKinney S, Thomas E. Evidence from multiplex molecular assays for complex multipathogen interactions in acute respiratory infections. J Clin Microbiol. 2008;46:97–102.

10. Peleg AY, Hogan DA, Mylonakis E. Medically important bacterial-fungal interactions. Nat Rev Microbiol. 2010;8:340–9. doi: 10.1038/nrmicro2313.

11. von Eiff C, Becker K, Machka K, Stammer H, Peters G. Nasal carriage as a source of Staphylococcus aureus bacteremia. Study Group. N Engl J Med. 2001;344:11–6.

12. Edouard S, Al-Tawfiq JA, Memish ZA, Yezli S, Gautret P. Impact of the Hajj on pneumococcal carriage and the effect of various pneumococcal vaccines. Vaccine. 2018;36:7415–22. doi: 10.1016/j.vaccine.2018.09.017.

13. Veenhoven R, Bogaert D, Uiterwaal C, Brouwer C, Kiezebrink HH, Bruin J, et al. Effect of conjugate pneumococcal vaccine followed by polysaccharide pneumococcal vaccine on recurrent acute otitis media: a randomised study. Lancet. 2003;361:2189–95. doi: 10.1016/S0140-6736(03)13772-5

14. Haut Conseil de la santé publique. Bull Epidemiol Hebdo. Recommandations sanitaires pour les voyageurs https://invs.santepubliquefrance.fr/content/.../62/file/Recommandations_voyageurs_2018.pdf; (2018)

15. Direction Générale de la Santé. Ministère des Affaires sociales, de la Santé et des Droits des femmes. Paris. Calendrier des vaccinations et recommandations vaccinales 2018, https://solidarites-sante.gouv.fr/IMG/pdf/calendrier_vaccinations_2018.pdf; (2018)

16. Al-Tawfiq JA, Gautret P, Memish ZA. Expected immunizations and health protection for Hajj and Umrah 2018-An overview. Travel Med Infect Dis 2017:19:2–7. doi: 10.1016/j.tmaid.2017.10.005.

17. Benkouiten S, Brouqui P, Gautret P. Non-pharmaceutical interventions for the prevention of RTIs during Hajj pilgrimage. Travel Med Infect Dis 2014:12:429–42. doi: 10.1016/j.tmaid.2014.06.005.

18. Hoang VT, Nguyen TT, Belhouchat K, Meftah M, Sow D, Benkouiten S, et al. Antibiotic use for respiratory infections among Hajj pilgrims: A cohort survey and review of the literature. Travel Med Infect Dis. 2019;30:39–45. doi: 10.1016/j.tmaid.2019.06.007.

19. Memish ZA, Assiri A, Turkestani A, Yezli S, Al Masri M, Charrel R, et al. Mass gathering and globalization of respiratory pathogens during the 2013 Hajj. Clin Microbiol Infect. 2015;21:571.e1–8. https://doi: 10.1016/j.cmi.2015.02.008.

20. Hoang VT, Meftah M, Anh Ly TD, Drali T, Yezli S, Alotaibi B, et al. Bacterial respiratory carriage in French Hajj pilgrims and the effect of pneumococcal vaccine and other individual preventive measures: A prospective cohort survey. Travel Med Infect Dis. 2018. pii: S1477–8939(18)30385-5. doi: 10.1016/j.tmaid.2018.10.021.

21. Memish ZA, Al-Tawfiq JA, Almasri M, Akkad N, Yezli S, Turkestani A, et al. A cohort study of the impact and acquisition of nasopharyngeal carriage of *Streptococcus pneumoniae* during the Hajj. Travel Med Infect Dis. 2016;14:242–7. doi: 10.1016/j.tmaid.2016.05.001.

22. Memish ZA, Assiri A, Almasri M, Alhakeem RF, Turkestani A, Al Rabeeah AA, et al. Impact of the Hajj on pneumococcal transmission. Clin Microbiol Infect. 2015;21:77.e11–8. doi: 10.1016/j.cmi.2014.07.005.

23. Hanada S, Pirzadeh M, Carver KY, Deng JC. Respiratory Viral Infection-Induced Microbiome Alterations and Secondary Bacterial Pneumonia. Front Immunol. 2018;9:2640. doi: 10.3389/fimmu.2018.02640.

24. Bischoff WE, Bassetti S, Bassetti-Wyss BA, Wallis ML, Tucker BK, Reboussin BA, et al. Airborne dispersal as a novel transmission route of coagulase-negative staphylococci: interaction between coagulase-negative staphylococci and rhinovirus infection. Infect Control Hosp Epidemiol. 2004;25:504–11.

25. Bischoff WE, Wallis ML, Tucker BK, Reboussin BA, Pfaller MA, Hayden FG, et al. “Gesundheit!” sneezing, common colds, allergies, and *Staphylococcus aureus* dispersion. J Infect Dis. 2006;194:1119–26.

26. Bischoff WE, Tucker BK, Wallis ML, Reboussin BA, Pfaller MA, Hayden FG, et al. Preventing the airborne spread of *Staphylococcus aureus* by persons with the common cold: effect of surgical scrubs, gowns, and masks. Infect Control Hosp Epidemiol. 2007;28:1148–54.

27. Bassetti S, Bischoff WE, Walter M, Bassetti-Wyss BA, Mason L, Reboussin BA, et al. Dispersal of *Staphylococcus aureus* into the air associated with a rhinovirus infection. Infect Control Hosp Epidemiol. 2005;26:196–203.

28. Merler S, Poletti P, Ajelli M, Caprile B, Manfredi P. Coinfection can trigger multiple pandemic waves. J Theor Biol. 2008;254:499–507.

29. Hofstra JJ, Matamoros S, van de Pol MA, de Wever B, Tanck MW, Wendt-Knol H, et al. Changes in microbiota during experimental human Rhinovirus infection. BMC Infect Dis. 2015;15:336. doi: 10.1186/s12879-015-1081-y.

30. Morgene MF, Botelho-Nevers E, Grattard F, Pillet S, Berthelot P, Pozzetto B, et al. *Staphylococcus aureus* colonization and non-influenza respiratory viruses: Interactions and synergism mechanisms. Virulence. 2018;9:1354–63. doi: 10.1080/21505594.2018.1504561.

31. Dunne EM, Murad C, Sudigdoadi S, Fadlyana E, Tarigan R, Indriyani SAK, et al. Carriage of *Streptococcus pneumoniae, Haemophilus influenzae, Moraxella catarrhalis*, and *Staphylococcus aureus* in Indonesian children: A cross-sectional study. PLoS One. 2018;13:e0195098. doi: 10.1371/journal.pone.0195098. eCollection 2018.

32. Pettigrew MM, Gent JF, Revai K, Patel JA, Chonmaitree T. Microbial interactions during upper respiratory tract infections. Emerg Infect Dis. 2008;14:1584–91. doi: 10.3201/eid1410.080119.

33. Xu Q, Almudervar A, Casey JR, Pichichero ME. Nasopharyngeal bacterial interactions in children. Emerg Infect Dis. 2012;18:1738–45. doi: 10.3201/eid1811.111904.

34. Ruohola A, Pettigrew MM, Lindholm L, Jalava J, Räisänen KS, Vainionpää R, et al. Bacterial and viral interactions within the nasopharynx contribute to the risk of acute otitis media. J Infect. 2013;66:247–54. doi: 10.1016/j.jinf.2012.12.002.

35. Perez AC, Pang B, King LB, Tan L, Murrah KA, Reimche JL, et al. Residence of *Streptococcus pneumoniae* and *Moraxella catarrhalis* within polymicrobial biofilm promotes antibiotic resistance and bacterial persistence in vivo. Pathog Dis. 2014;0:280–8. https://doi.org/10.1111/2049-632X.12129 PMID: 24391058

36. Weimer KED, Juneau RA, Murrah KA, Pang B, Armbruster CE, Richardson SH, et al. Divergent mechanisms for passive pneumococcal resistance to β-Lactam antibiotics in the presence of *Haemophilus influenzae*. J Infect Dis. 2011;203:549–55. https://doi.org/10.1093/infdis/jiq087.

37. Direction Générale de la Santé. Ministère des Affaires sociales, de la Santé et des Droits des femmes. Paris. Calendrier des vaccinations et recommandations vaccinales 2019, https://solidarites-sante.gouv.fr/IMG/pdf/calendrier_vaccinal_mars_2019.pdf; (2019)

38. Rashid H, et al. Influenza and the Hajj: defining influenza-like illness clinically. Int J Infect Dis. 2008;12:102–3.

39. WHO. Body mass index – BMI http://www.euro.who.int/en/health-topics/disease-prevention/nutrition/a-healthy-lifestyle/body-mass-index-bmi; (2019)

40. Ninove L, Nougairede A, Gazin C, Thirion L, Delogu I, Zandotti C, et al. RNA and DNA bacteriophages as molecular diagnosis controls in clinical virology: a comprehensive study of more than 45,000 routine PCR tests. PLoS One. 2011;6:e16142

41. Greiner O, Day PJ, Altwegg M, Nadal D. Quantitative detection of *Moraxella catarrhalis* in nasopharyngeal secretions by real-time PCR. J Clin Microbiol. 2003;41:1386–90.

